# Neural correlates of appetitive extinction learning: An fMRI study with actively participating pigeons

**DOI:** 10.1101/2025.04.28.650993

**Authors:** Alaleh Sadraee, Xavier Helluy, Erhan Genç, Meng Gao, Mehdi Behroozi, Onur Güntürkün

**Affiliations:** Department of Biopsychology, Faculty of Psychology, Institute of Cognitive Neuroscience, Ruhr University Bochum, Universitätsstraße 150, 44780 Bochum, Germany; International Graduate School of Neuroscience, Ruhr University Bochum, Bochum, Germany; Department of Neurophysiology, Faculty of Medicine, Ruhr University Bochum, Universitätsstraße 150, 44780 Bochum, Germany; Department of Psychology and Neuroscience, Leibniz Research Centre for Working Environment and Human Factors (IfADo), Ardeystraße 67, 44139 Dortmund, Germany; Developmental Computational Psychiatry lab, Department of Psychiatry and Psychotherapy, Faculty of Medicine, University Tübingen, Calwerstraße 14, 72076 Tübingen, Germany

**Keywords:** Bird, Prefrontal, Hippocampus, Go/NoGo Paradigm, Brain Asymmetry, Multiple Memory Systems

## Abstract

Extinction learning is an important learning process that enables adaptive and flexible behavior. Human neuroimaging studies show that the neural basis of extinction learning consists of a neural network that includes the hippocampus, amygdala, and subcomponents of the prefrontal cortex, but also extends beyond them. The limitations of applying fMRI to actively participating animals have so far restricted the identification of the entire extinction network in non-human animals. Here, we present the first fMRI study of extinction in awake and actively participating pigeons, using a Go/NoGo operant paradigm with a water reward. Our study revealed an extensive and largely left hemispheric telencephalic network of sensory, limbic, executive, and motor areas that slowly ceased to be active during the process of extinction learning. We propose that the onset of extinction ignites a neuronal updating of the associated consequences of own actions within a large telencephalic neural network until a new association is established which inhibits the previously acquired operant response.

## Introduction

Animals must flexibly adapt their behavior to the ever-changing conditions of their surroundings. Thus, both the ability to learn and the aptitude to subsequently extinguish the association between a conditioned (CS) and an unconditioned stimulus (UCS) is of key importance. Extinction does not largely erase the CS-US acquisition memory^1^ but establishes a second inhibitory trace that suppresses the occurrence of the conditioned response (CR)^2^. Most studies on the neural mechanism of extinction learning were conducted in rodents using aversive Pavlovian conditioning^3^. These investigations point to the involvement of the amygdala, prefrontal cortex (PFC), and hippocampus as key entities of aversive extinction learning^3,4^. While this seems to apply for conditioned fear extinction in rodents, studies in human participants note some differences like for the amygdala^5^. Some studies therefore assume that the amygdala quickly adapts over time within a few trials while other studies cannot find evidence for this claim^6,7^. In addition, aversive and appetitive learning seems to involve partly different neural mechanisms^8–10^. Many further studies that used various conditioning procedure also discovered a large variety of cortical and subcortical areas that also accompanied extinction learning^11–16^. Thus, a variety of structures participate in extinction.

Extinction learning is widespread within the animal kingdom and has also been demonstrated in many non-mammalian species like birds^17^, fish^18^, and invertebrate species as honeybees^19^ and fruit flies^20^. To our knowledge, a systematic search for the neural fundaments of appetitive extinction in vertebrates outside the mammalian class has only been conducted in pigeons. This was done by training these animals in a within-subject renewal design in which pigeons were classically conditioned to peck in a context-dependent manner on two conditioned stimuli. Before extinction learning onset, birds received local and area-specific injections of saline or drug (either AP5 or TTX). These studies revealed that, like in mammals, hippocampus, amygdala, and the nidopallium caudolaterale (NCL), the avian functional equivalent of the prefrontal cortex^21,22^, play key roles in different aspects of extinction learning. In addition, blocking the visual associative nidopallium frontolaterale (NFL)^23^ slowed down extinction and impaired retrieval of context-specific memory^24^. Inhibiting NMDA receptors in the striatum and premotor arcopallium reduced extinction learning speed^24–26^ and extinction memory consolidation, respectively^26^. To summarize, the visual-associative NFL, the ‘prefrontal’ NCL, as well amygdala and striatum participate in appetitive extinction learning in birds^27^. Hippocampus, NCL, and premotor arcopallium contribute to extinction memory consolidation, while NFL and NCL play key roles in the contextual embedding of extinction learning^24^. Such a pattern overlaps to some extent with results in mammals^3^ but also shows some differences. Since the evolutionary lines of extant mammals and birds parted 324 million years ago^28^, a rudimentary telencephalic network for extinction learning seems to have existed since the early times of land vertebrates and subsequently accumulated some mammalian- or avian-specific systems.

All of the above cited studies in pigeons targeted one area and described the subsequent behavior as a one-shot change. To identify all telencephalic network changes throughout the process of extinction, a completely different approach is needed. This is the reason why we decided to use fMRI in awake and actively participating pigeons. fMRI provides a non-invasive, system-wide measurement of neuronal activation with a decent temporal and high spatial resolution. In the last years, we established a robust fMRI protocol that has been tailored for use with awake, behaving pigeons^29^. To study extinction learning at whole system levels with fMRI, we switched our learning paradigm to an operant Go/NoGo design with water as a reward. This is different from the Pavlovian design in the above-mentioned studies and might go along with differences in the involved areas. To our knowledge, this is the first fMRI study on extinction learning in a non-human animal.

## Results

Behavioral and neural data were recorded inside the 7T small animal MRI scanner while pigeons went through an appetitive operant discrimination paradigm, followed by extinction the next day (24h later). Briefly, 8 pigeons were first surgically implanted with an MRI-compatible pedestal that was later used to fix to a restrainer and keep their head fixed during experimental sessions (Figure 1a). Animals then went through a three-step habituation procedure to be accustomed to our imaging protocol. Finally, water-deprived pigeons were trained for a Go/NoGo task as an appetitive operant discrimination paradigm with water as a reward and mandibulation (lower jaw opening) as an operant response. As a result, pigeons learned to discriminate between two colored cues and to mandibulate only after the appearance of the Go cue (red or green color, S+) and to inhibit responses during the NoGo cue (green or red color, S-). Behroozi et al. (2020)^29^ presented brain activations during discrimination between these two cues.

**Fig 1.**
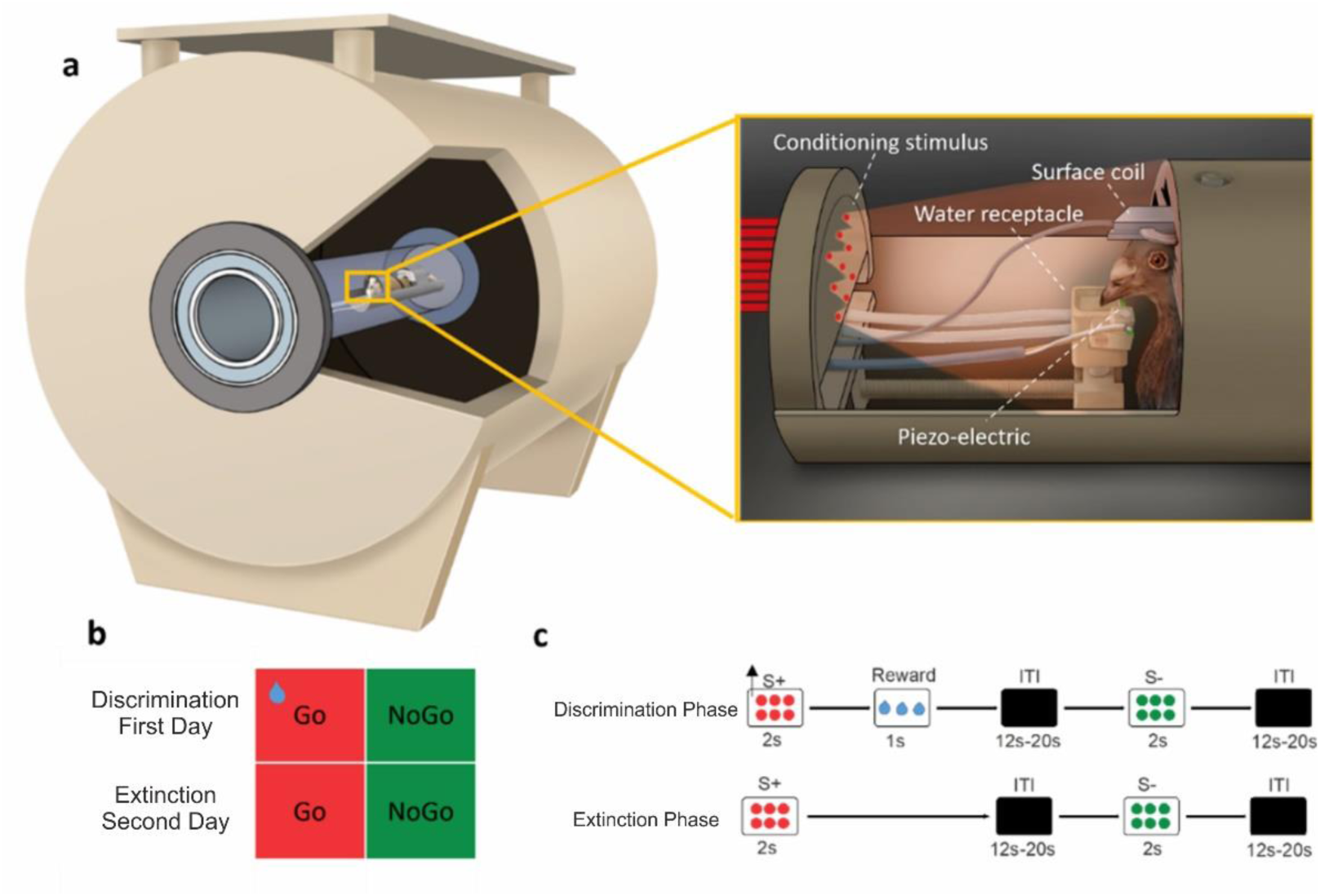
Experimental set-up and task design for behaving pigeons inside the MRI scanner. **(a**) Custom-made restrainer and 7T small animal MRI device. This system was designed to immobilize the animal by fixing its head and body. Fiber optics for visual stimulus presentation were placed in front of the pigeon. A custom-made light emitter was designed and placed outside the scanner room. A piezo-electric device was positioned under the lower jaw to register the animals’ responses. To reward pigeons after a correct response, a small receptacle was placed at the tip of the beak, which could be filled with water. (**b)** The experiment was performed on two consecutive days, with the discrimination task conducted on the first day, followed by extinction on the second. The task consisted of two types of trials: Go (S+) and NoGo (S-) trials **(c)** visual stimuli were presented for 2s, which were followed with jittered ITI (12.2-20.2 s). During the discrimination experiment, a water reward was delivered after responding to the S+, while no reward was provided during extinction. In both phases, S-trials were not rewarded, regardless of the presence or absence of operant responses. Arrows represent animal mandibulations which result in water reward (blue drop).

Twenty-four hours after a successful acquisition of the color discrimination, pigeons underwent extinction by withholding appetitive water reward after a response to the Go cue, while BOLD brain responses were simultaneously recorded (Figure 1b, c). Each fMRI recording session consisted of two blocks, each containing 72 trials (36 Go and 36 NoGo), separated by a 20-minute resting period. For the subsequent analysis, the first block will be referred to as the early phase of extinction, while the second block will be termed the late phase of extinction.

### Behavioral responses during discrimination and extinction

The fractional mandibulation rate during S+ and S-presentation for 2s was separately calculated for each pigeon. During the discrimination phase, the fractional mandibulation rate for S+ (0.77 ± 0.03; mean ± SEM) was significantly higher than for S- (0.14 ± 0.02) (two-tailed paired t-test, t_(7)_ = 17.27; *p* < 0.001, Figure 2a). Throughout the extinction phase, the fractional mandibulation rate for S+ (0.43 ± 0.06; mean ± SEM) remained significantly higher than for S- (0.10 ± 0.03) (two-tailed paired t-test, t_(7)_ = 6.85; *p* < 0.001, Figure 2a), but it significantly decreased from the discrimination phase to the extinction phase for S+ (two-tailed paired t-test, t_(7)_ = 5.25; *p* = 0.001).

**Fig 2.**
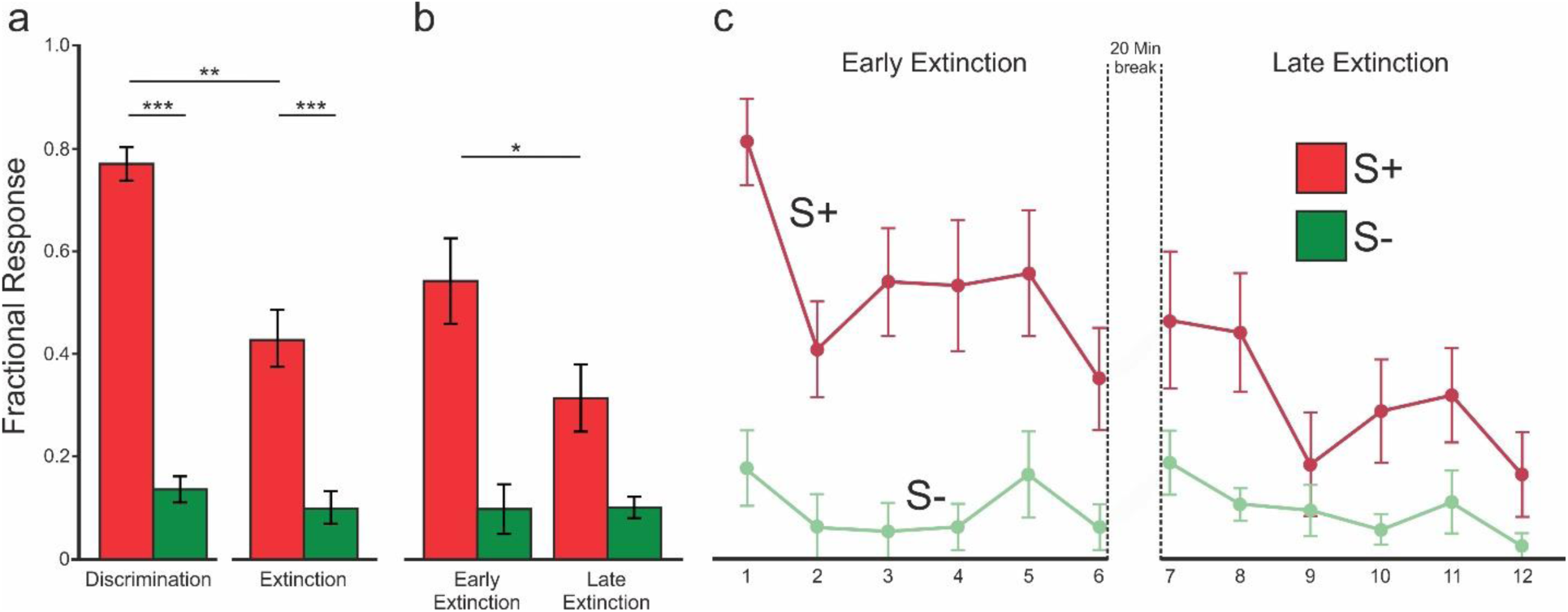
Behavioral Results. a. Behavioral results of the entire discrimination (day 1) and entire extinction phase (day 2); The mean fractional mandibulation rate for Go (S+, red) and NoGo (S-, green) differ significantly between S+ and S-stimuli in both phases. b. The extinction phase is segmented into early and late phases; findings indicate a time-related reduction of mandibulation rate for S+, while responses to the S-remain unchanged. c. The plot shows the gradual decline in responding to S+ during extinction; The data are grouped into bins of six consecutive trial for both S+ and S-. Error bars indicate the standard error of the mean (SEM). * p= 0.06; ** p< 0.05; *** p< 0.001.

To investigate the effect of stimulus type (S+ vs. S-) and phase (early vs. late extinction; defined by first and second block, respectively), we performed a two-way ANOVA. The result indicated a statistically significant effect of stimulus type (F_(1, 7)_ = 46.94; *p* <0.001), no main effect of phase (F_(1, 7)_ = 3.07; *p* = 0.12), and a significant interaction between stimuli type and phase (F_(1, 7)_ = 6.66; *p* = 0.036) suggesting that the response to S+ and S− stimuli differed across the early and late phases of extinction.

To assess how the response to S+ and S− changed across extinction phases, the fractional mandibulation rate for both S+ and S-was evaluated across time. This revealed a strong trend toward a time-dependent decrease of responses to the S+ (two-tailed paired t-test, t_(7)_ = 2.2; *p* = 0.06, Figure 2b), while responses to the S-remained unaltered (two-tailed paired t-test, t_(7)_ = 0.1; *p* = 0.93). Therefore, we analyzed the fMRI contrasts separately for the first and the second halves of extinction phases.

To illustrate the trend of response during extinction, fractional responses for both stimulus types were grouped into bins of twelve consecutive trials. A gradual decline in responding during extinction was observed for S+, while the trend for S-remained largely unchanged (Figure 2c).

### fMRI results

To identify the brain activity during the early and late phases of extinction, we analyzed the BOLD fMRI contrast S+_ext_ > S-_ext_ for the first and second blocks of the extinction phase. S+_ext_ was defined as the Go trials during extinction in which the birds did not respond. Accordingly, S-_ext_ referred to NoGo trials during extinction in which the animals also did not respond. See the method section for more details regarding the choice of this contrast.

First-level analyses were conducted in single-subject space using the contrasts S+_extE_> S-_extE_ (early phase) and S+_extL_ > S-_extL_ (late phase). The resulting parameter estimates (beta values) for each subject were entered into the second-level group analysis using random-effect modeling. Statistical thresholds were set at Z = 3.1, with a p < 0.05 family-wise error (FWE) correction applied at the cluster level.

### Early extinction phase

During early extinction phase, overall, seven activity clusters were identified. They are depicted in the small insets of figure 3 in different colors. Four of them were identified in the right hemisphere. These included activations localized to the hippocampus (purple), the striatum (blue), the MIVl (light pink), and a larger cluster in the right hemisphere encompassing multiple sensory and sensory-associative regions (E, MFV, MID, MIVl, MIVm, NFL, NIL, NIMl, NMm; green). In the left hemisphere, significant clustered activation was observed in the TuO (dark pink), and NCC (gray). A larger cluster (red) also exhibited significant activation, encompassing sensory-associative (E, HA, HD, HI, IHA, MIVl, NIL, TPO), limbic (MFD, NCC, NDB, NSTL), and sensorimotor (AD, MIVm, NIMl, NMm, StM, StL, VP) areas.

**Fig 3.**
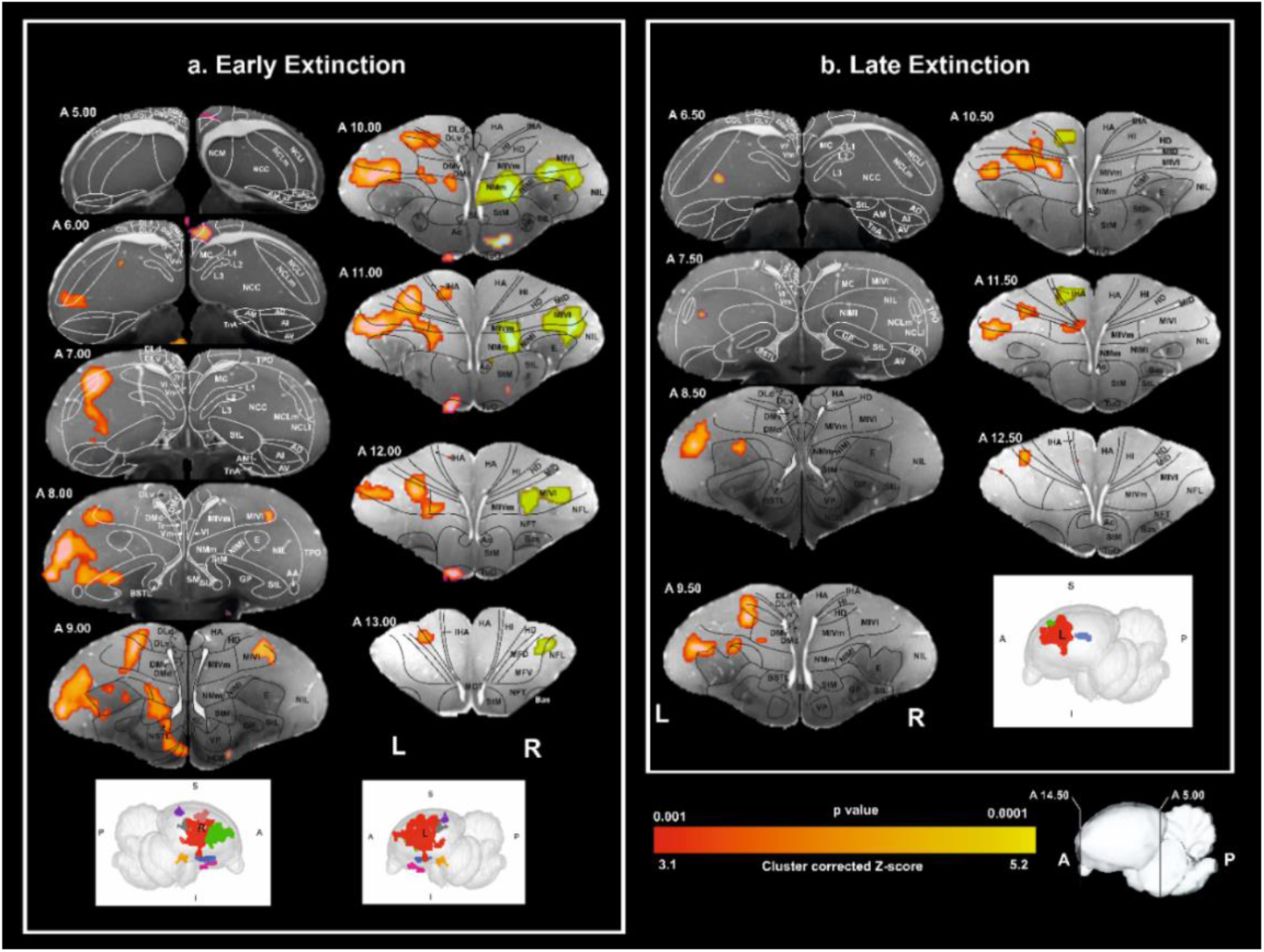
Activation patterns during extinction phases. Coronal images at different levels of the pigeon telencephalon were registered onto a pigeon brain MRI atlas^52^. Significantly activated areas with a cluster threshold of Z> 3.1 and FWE p<0.05 are highlighted in color. BOLD fMRI contrasts S+_ext_ > S-_ext_ were analyzed for the early (first block) and late (second block) extinction phases. S+_ext_ refers to Go trials during extinction with no response, and S-_ext_ refers to NoGo trials with no response. Early Extinction Phase (a): Right Hemisphere: Hippocampus (purple), Striatum (blue), MIVl (light pink), and a larger cluster (green) in sensory and sensory-associative regions. Left Hemisphere: TuO (dark pink), NCC (gray), and a larger cluster (red) in sensory-associative, limbic, and sensorimotor areas. Late Extinction Phase (b): Left Hemisphere: NCC (blue), HA/IHA (green), and a larger cluster (red) in sensory and sensory-associative structures. For anatomical abbreviations see Table 1.

**Table 1:**
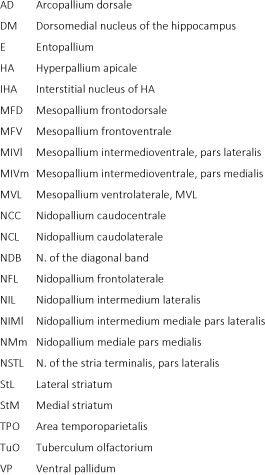
List of Anatomical Abbreviations AD Arcopallium dorsale.

### Primary and secondary sensory areas

Starting anterior at A13.50 and leading until the posterior end of the visual wulst, the left hyperpallium intercalatum (HI) and the hyperpallium densocellulare (HD), two thalamopallial input areas of the visual thalamofugal pathway evinced significant BOLD responses^30^. The avian thalamofugal system corresponds to the mammalian geniculocortical visual system^31,32^. Beginning with A11.50 and A11.00, also the hyperpallium apicale (HA) and the interstitial nucleus of HA (IHA) were activated respectively. While the HA receives secondary input from all wulst layers^33^, IHA receives the bulk of the thalamopallial visual input from the N. geniculatus lateralis pars dorsalis^34^. No corresponding activation was seen in the right hemisphere.

In addition to the activation of the thalamofugal pathway, also parts of the left, but not the right entopallium were activated. The entopallium is the thalamopallial projection area of the n. rotundus of the avian tectofugal visual system which corresponds to the mammalian extrageniculocortical pathway^32^. In addition to the primary visual entopallium, also secondary visual tectofugal areas were bilaterally activated. These are from anterior to posterior, Nidopallium frontolaterale (NFL), Nidopallium intermedium lateralis (NIL), and Mesopallium intermedioventrale, pars lateralis (MIVl, incl. the Mesopallium ventrolaterale, MVL), and^35,36^. Widespread BOLD signals were also detected in the left hemispheric Area temporoparietalis (TPO) which receives input from pallial structures of both the tecto- and the thalamofugal system^37^ and is involved in form, color, and motion discriminations^23^.

The Mesopallium frontoventrale (MFV) was activated in the right hemisphere. The MFV is a higher associative area of the trigeminal system and has reciprocal projections to primary and secondary pallial trigeminal areas, the “prefrontal” NCL and the premotor arcopallium^35^. We also detected BOLD signals in the Tuberculum olfactorium (TuO) on the left side. The TuO is, like in mammals, conceivably the recipient of olfactory bulb input^38^ (but see^39^). Finally, BOLD signals were bilaterally present in the Mesopallium intermedioventrale, pars medialis (MIVm), a multimodal association area^35^.

### Limbic areas

The Nidopallium caudocentrale (NCC), a higher-order limbic forebrain area^35^, revealed left hemispheric BOLD signals in its lateral components. We also detected BOLD responses in the Mesopallium frontodorsale (MFD), a limbic association area that reciprocates with the amygdala and hippocampal formation^35^. Finally, as part of the extended amygdala, the N. of the stria terminalis, pars lateralis (NSTL) was active, as also the the N. of the diagonal band (NDB) that has reciprocal interactions with the hippocampus^40^.

### Executive control

We evinced a left hemispheric activation within the borders of the “prefrontal” NCL^41^. Comparing the fields of NCL-activations with the anatomical reconstruction of the secondary sensory afferents of this structure^42^, it became evident that the BOLD signals were confined to those parts that integrated trigeminal and visual tectofugal and thalamofugal input. The hippocampus of the right hemisphere was activated at levels A 5.0 and A 6.0. BOLD signals were confined to the dorsomedial hippocampal component, encompassing both the dorsal (DMd) and the ventral (DMv) part^43^.

### Response production

The Arcopallium dorsale (AD) was activated between A7.00 and 6.50, with the field of activity moving dorsally into the NCL in more posterior sections. The AD is a premotor area that nevertheless also receives input from several limbic structures^44^. In more anterior sections, BOLD signals were detected along the medial wall of the mesopallium and nidopallium. The three pallial nuclei in this region, the Mesopallium intermedioventrale, pars medialis (MIVm), the Nidopallium intermedium mediale pars lateralis (NIMl) and the Nidopallium mediale pars medialis (NMm) are involved in fast sensorimotor learning^45,46^ and sequential behaviour^47–51^. They all showed bilateral BOLD signals during early extinction learning. At the subpallial level, the left hemisphere evinced activations of the lateral (StL) and the medial striatum (StM) as well as in ventral pallidum (VP).

### Late extinction phase

During late extinction phase, only three left hemispheric activity clusters were identified (Fig. 3, 4). Two small ones were isolated to the NCC (blue) and HA/IHA (green), while a larger cluster (red) encompassed regions including sensory and sensory-associative structures (E, HD, HI, MFD, MIVl, MIVm, NIL, and NIMl). In the primary and secondary sensory areas, all left hemispheric thalamugal visual areas (HA, IHA, HI, HD) evinced activity patterns that extended along the full length of the visual wulst, although the extent of BOLD signals was smaller. Also, the left primary visual entopallium (E) as well the tectofugal associative components NIL and MIVl were active. Again, the BOLD signal was considerably smaller but occupied a similar anterior-to-posterior extent. It is important to note that the cortex-like columnar system of the thalamofugal (HA, IHA, HI, HD) and the tectofugal systems (MIVl, NIL, E) were always active as a group, strengthening the assumption that they act as one canonical unit^33^.

**Fig. 4.**
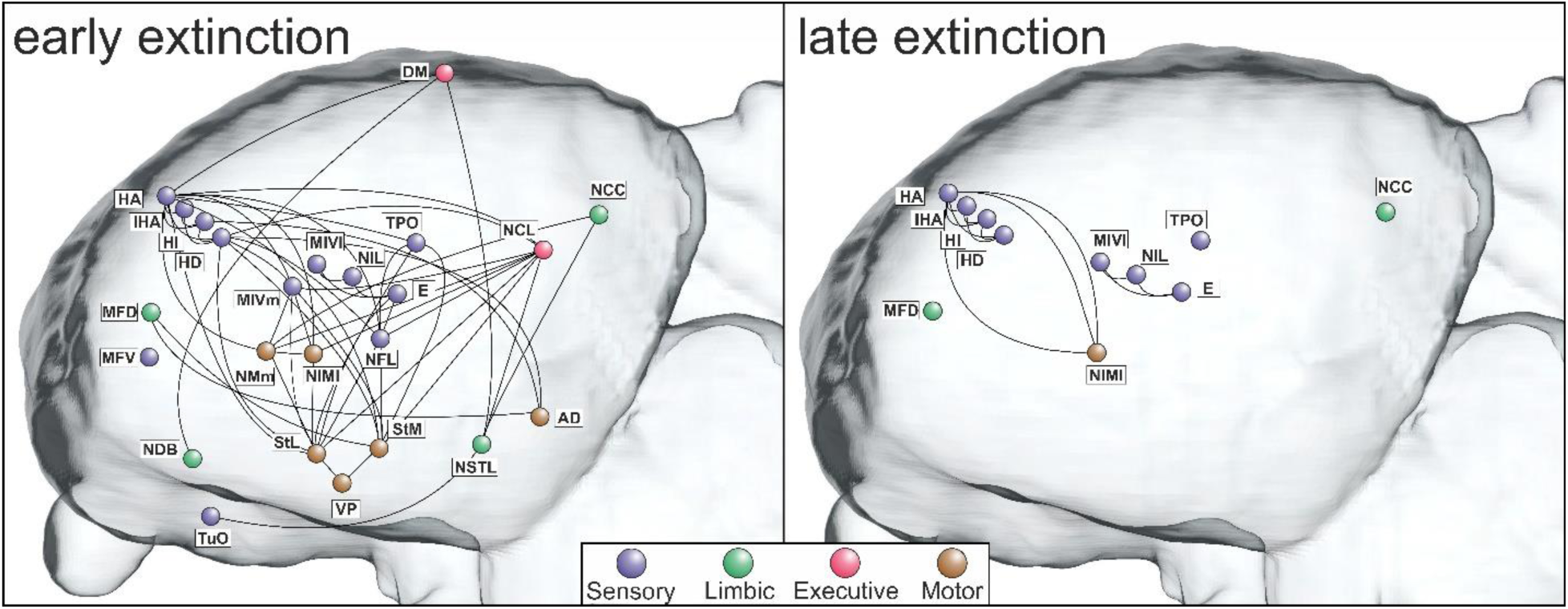
Network of activations during early and late extinction. Saggital sections of activated areas during early and late extinction phases, organized into four functional systems depicting sensory, limbic, executive, and motor areas. Left and right hemispheric results were collapsed. Connections are derived from tracing studies^36^. Note cortex-like columnar activations in thalamofugal (HA, IHA, HI, HD) and tectofugal visual areas (MIVl, NIL, E). For anatomical abbreviations see Table 1.

At mesopallial level, the MFD as the higher order limbic area showed a small area of activation. Of the pallial higher order motor areas, only the left sided MIVm and, to a very small extent, also the NIMl were active.

## Discussion

We conducted the current fMRI study to identify extinction learning dependent activity changes of the entire avian telencephalic network. Appetitive extinction studies in pigeons take longer than fear extinction experiments in rodents^53^. This gives us the opportunity to not only identify the extinction-related neural components, but to also reveal how activity patterns change over learning time. We therefore separately analyzed and compared the data set for the first and the second half of the extinction session. Overall, we departed from the view that memory is supported by multiple brain systems that are simultaneously active and are subsequently modified in parallel during extinction^54,55^. We therefore organized our discussion into processes that reflect stimulus-stimulus and stimulus-affect associations as well as executive control and response production.

### Stimulus-stimulus associations

Bolles^56^ was among the first who argued that associative learning does not only establish a conditioned S-R but also an S-S association. This effect is especially visible in appetitive operant tasks in which animals start approaching the expected location of reward delivery when perceiving the conditioned stimulus^57^. If this expectation turns out to be wrong during extinction, task contingencies have to be updated, resulting in major changes of neuronal firing patterns across brain areas^58^. Conceivably, these alterations go along with learning of a secondary association that suppresses the initial response to the conditioned stimulus^2,4^. We assume that most of the activity patterns that we observed, result from expectation updating and the subsequent forming of a second association of inhibitory nature^27^. For the sensory areas, updating was possibly related to the stimuli that accompanied the expected outcome of the operant response. During learning of the color discrimination, the animals associated a correct response with water reward. This reinforcement was embedded in a rich compound of sensory events. First, after responding to the S+, water reward was signaled by two white lights that activated the visual system. Second, the pigeons sucked and ingested the water – an action with kinesthetic trigeminal feedback^59^. Third, a gustatory and possibly an olfactory input accompanied drinking. If extinction goes along with a modification of brain systems that accompanied the conditioned response during acquisition, we should expect these areas to also be active during extinction.

During early extinction phase, we indeed found a massive activation of the relevant sensory and sensory-associative areas of the avian pallium. These included primary and secondary structures of the visual tectofugal (E, NFL, NIL, MIVl, TPO) and visual thalamofugal (HA, HD, HI, IHA) systems^32,60,61^. In addition, sensory-associative structures of the trigeminal (MFV) and the olfactory system (TuO) were active, as also the MIVm, an association area that integrates all senses and serves as a higher-order sensorimotor area^35,62^. In the late extinction phase, the BOLD signal in all of these areas shrank in size, while no significant activation was detected in TuO. Taken together, our data make it likely that despite our pigeons had stopped responding, updating of the expected sensory events resulted in BOLD responses across sensory areas. These diminished from the early to the late extinction phase.

### Stimulus-value associations

We did not see activity patterns in the amygdala, but in other limbic areas that closely interact with this structure. This is foremost the NCC, a region located in the central caudal nidopallium that in songbirds is involved in appetitive events like courtship^63^ and that interacts with subnuclei of the avian amygdala and the nucleus of the stria terminalis^64^. This latter structure was also activated during extinction learning. A further active limbic area during extinction was the limbic associative MFD that reciprocates with the amygdala and, via the Area corticoidea dorsolateralis (CDL), with the hippocampal formation^35^. Also, the N. of the diagonal band (NDB) was activated – an area that reciprocates with the hippocampal formation^40^. All these regions were less (NCC, MFD) or no longer active (NDB, NSTL) during late extinction.

### Executive control

The hippocampus plays a key role in stimulus-stimulus associations, especially when connecting cues across sensory domains^65^. During the early phase of extinction, the avian hippocampal DM was active - a central hub of the avian hippocampal system that is structurally and functionally strongly connected to the input structures of the hippocampal formation^43,66^. DM is also at the crossroad of two major transversal pathways that loop within the avian hippocampus^67^. Thus, DM could play a key role as a gatekeeper for sensory information that enters the hippocampus and could participate in neuronal updating of the associated consequences of own actions during extinction. Indeed, Lengersdorf et al.^17^ discovered that a pharmacological blockade of the pigeon hippocampus impedes extinction learning. This accords with fMRI studies in humans that report hippocampal activations during the early extinction phase^68,69^.

Extinction learning also activated the avian nidopallium caudolaterale (NCL) which is an associative pallial area that is functionally equivalent to the prefrontal cortex^70^. Like the mammalian prefrontal cortex, the avian NCL is also strongly innervated by dopaminergic midbrain neurons^21,71^ and displays reward prediction errors during extinction learning^27,72^. Lissek & Güntürkün^73^ and Lengersdorf et al.^25^ showed that local inhibition of the NMDA receptors in the NCL slowed down acquisition of extinction, while transiently inactivating the NCL with the Na+ channel blocker tetrodotoxin (TTX) impaired extinction memory consolidation^17^. This is like various extinction studies in mammals, incl. humans^12,13,58,74^.

Stimulus-response associations: While the subpallial components of the avian motor system are to some extent known^75,76^, studies on the relevant pallial structures are more limited. It is known that the arcopallium serves as a premotor structure^44^ with some of its descending projections to brainstem motor nuclei activating beak movememts^77^. Recordings from arcopallial neurons show that many of them start firing after perceiving the S+ and stop after response execution^78^. Thus, the AD that was activated in the current study, represents an essential link of the avian sensorimotor system^49^ and also integrates input from limbic areas^79^. In more anterior sections, three small structures (MIVm, NIMl, NMm) along the medial wall of the mesopallium and nidopallium were active. These receive multimodal input, reciprocate with NCL and arcopallium and have projections to the striatum^35,36,80^. They play a role in fast sensorimotor learning and, together with the NCL, in sequential behaviour^45,49–51^. Deactivation of NIML goes along with an increase of reaction times and sequence specific errors in serial response tasks, indicating that NIML could process a ‘‘what comes next’’ function^47,48,50^. The instrumental Go/NoGo-task with mandibulation as the operant was learned by our pigeons in a lengthy training that possibly established, at least in part, a habit-based response to the S+^81^. The goal-directed system in both rats and humans depends on the posterior dorsomedial striatum^82,83^, while the habit system is mostly governed by the dorsolateral striatum^84,85^. Corresponding information is missing in birds, but we identified significant BOLD responses in both medial and lateral striatal areas (StM, StL) as well as in the globus pallidus (GP). This makes a partly habit-based response pattern conceivable. Indeed, Gao et al.^24^ showed that blocking NMDA of the StM substantially deteriorated extinction acquisition in pigeons.

Throughout extinction, visual areas of the left hemisphere were more active than those of the right. Although this asymmetry was now for the first time detected with fMRI, it is well known that discrimination of color and further features of visual objects^86,87^ relies in many bird species on a superiority of left hemispheric circuits^88–90^. In pigeons, the asymmetry of visual feature discrimination arises through an interaction of genetic and epigenetic factors during embryonic age^91^, alters the animals pallial neural circuitry^92,93^ and can be detected both during natural behavior like homing^94^ and when tested during conditioning studies^95^.

Studies on the neuronal basis of this asymmetry could show, that left hemispheric visual tectofugal neurons develop higher activity bursts when visual features are associated with reward^96^. In addition, the activated neurons also show a better differentiation between rewarded and unrewarded cues^96^. This last point implies that during choice situations, left hemispheric visual neurons become better predictors of future rewards, thereby strengthening the synaptic coupling between left hemispheric areas that reach from visual via executive to motor structures^97^. It is important to remind that the current task did not test acquisition but extinction. The current discovery of an extinction-associated lateralized activation of a left hemispheric network could imply that both acquisition and extinction rests on overlapping left hemispheric structures.

The current study demonstrated that extinction learning in pigeons goes along with widespread activity patterns in sensory, limbic, executive, and motor areas. These activations were mostly left hemisphere-based and ceased during the progress of extinction learning. The classic notion of fear extinction studies in rodents posits that especially parts of the prefrontal cortex, the hippocampus as well as the amygdala constitute core components of the extinction network. Our findings only party overlap with this assumption but accords with the view that different learning procedures (aversive - appetitive; Pavlovian – instrumental) and species (rats/mice – pigeons) result in different neural representations that are subsequently updated when prediction errors ignite extinction^98^. Thus, our current data conceivably describe the areal distribution of these neural changes during extinction learning^29,55,99^.

Packheiser et al.^27^ recorded from single NCL neurons during extinction learning and discovered that the reward prediction error was ignited as a peak of population activity by reward omission. It then moved backwards in time towards trials that represented the presentation of the S+, to then move further to those trials that coincided with changes of decision-making. Importantly, this reward prediction error was expectancy-driven because it was elicited by changes in reward expectation, even when neither the quality nor quantity of reward was altered. Thus, the multimodal and associative neurons of the NCL changed their activity patterns that coded for reward expectations, associated sensory signals, and ensuing actions.

While the experiment of Packheiser et al.^27^ was focused on the NCL, the current study could analyze the whole telencephalic network. We uncovered a rich network of sensory, limbic, executive, and motor areas that accompanied extinction. This accords with data from Russo et al.^58^ who recorded from the prelimbic PFC of rats that underwent extinction learning of an operant response. The authors observed a population-wide neuronal transition during updating of task contingencies, such that ceasing of the operant response went along with activities in task-activated neural networks that continuously interacted with neurons that command dissimilar sub-functions. Like our fMRI-based observation, Russo et al.^58^ observed that both acquisition and subsequent extinction are constituted by overlapping and transiently activated interconnected neural systems. We assume that this also applies to pigeons and that the activated telencephalic networks participate in the acquisition of a new association of inhibitory nature that then stops the operant response.

Overall, our data show a much wider and more diverse activated network than often reported during extinction studies. Does this imply that the often-used term “extinction system” suggests an entity that in a strict sense is questionable? If associative learning takes place in different species and involves different principles (Pavlovian, instrumental), response patterns, sensory cues, and cognitive components, we might expect varieties of brain systems to undergo synaptic modification during extinction. But it is conceivable that a small cluster of structures is always involved, regardless of the experimental variations listed above? Indeed, our study identified prefrontal and hippocampal areas to be activated during extinction. Although we found no BOLD responses in the avian amygdala, other limbic structures were active. Thus, extinction learning might go along with an activation of an invariant core network of executive, memory-related, and limbic structures. To test this hypothesis, we need experimental approaches that enable whole system analyses combined with variations of extinction learning designs. We hope that our study contributed to this aim.

## Materials and Methods

### Subjects

Eight adult pigeons (Columba livia figurita) obtained from local breeders served as subjects (weight: 150–200 g, 3 years old, undetermined sex). Birds were individually housed in wire-mesh cages inside a temperature- and humidity-controlled colony room under a 12-h light/dark cycle with food and water ad libitum during the recovery period. During the experimental phase, pigeons were deprived of water for 12h before the start of the experiment to increase their motivation to be engaged in the experimental procedures.

### Ethical Statement

All animals were treated in accordance with the German guidelines for the care and use of animals in science. In addition, all procedures were reviewed and approved by the national ethics committee of the State of North Rhine-Westphalia, Germany (Landesamt für Natur, Umwelt und Verbraucherschutz Nordrhein-Westfalen (LANUV), Application number: Az.:84– 02.04.2014.A206). These procedures are in accordance with the ARRIVE guidelines (https://arriveguidelines.org).

### Surgery

Each pigeon was surgically implanted with an MR-compatible plastic pedestal attached to the skull. The procedure has already been published by Behroozi et al., 2020. In summary, before surgery, animals were anesthetized with a mixture of Ketamine (100 g/m; Pfizer GmbH, Berlin Germany) and Xylazine (20 mg/ml Rompun, Bayer Vital GmbH, Leverkusen Germany) in a 7:3 ratio (0.075ml/100gr). In addition, gas anesthesia (Isoflurane, Forane 100% (V/V), Mark 5, Medical Developments International, Abbott GmbH & Co KG, Wiesbaden, Germany) has also flowed through a mask for assured anesthetization through the surgery process. The animals were then placed in a stereotactic apparatus. An incision was made to expose the skull. After cleaning the skull, four Polyether ether ketone (PEEK) micro pan head screws were placed in the anterior and posterior surface of the skull. A custom-made plastic pedestal was attached to screws with dental cement (Omni Ceram). Dental cement is then used to seal the surrounding of the pedestal to increase the adhesive strength between the skull and the pedestal. After surgery, analgesics (Carprofen (Rimadyl), 10 mg/kg) and antibiotic (Baytril, 2.5 mg/kg) were applied twice daily on three consecutive days. Animals were allowed to recover for at least 6–8 weeks before habituation started.

### Habituation

Pigeons were habituated to the MRI imaging environment inside a mock scanner, mimicking the real MRI environment located in a habituation room (outside the scanner). We used a well-established protocol in our laboratory proven to minimize body motion and stress associated with head fixation^66,100^. Briefly, in three consecutive phases, pigeons were first wrapped in a cloth jacket and placed in the restrainer inside the mock scanner setup for three days. Then, they were fixed to the restrainer via the implanted plastic pedestal on their skull for the time started from 10 min and were increased stepwise up to 100 min. Finally, a recording of the real scanner noise was introduced to the animals, gradually increasing up to 90 dB (the same sound pressure as that inside an actual animal scanner).

### Apparatus

The platform to run behavioral experiments has already been established by Behroozi et al.^29^. It consists of four main modules: (i) a mock scanner with the holding device used for maintaining the animal in a head-fixated state; (ii) a piezo-electric sensor placed under the lower mandible for operant response registration; (iii) a water receptacle, positioned at the tip of the beak that allowed animals to access the water reward upon their correct responses; water delivery was controlled by two peristaltic pumps (SR 10_50 12V DC Tubing Novoprene) that were connected to a water supply reservoir; (iv) a 3×3 LED matrix (TLC5947 PWM LED Driver) for stimulus presentation. The stimuli were transferred inside the scanner using non-magnetic fiber-optics. All hardware of the setup was controlled with a digital input/output interface (Arduino Mega 2560), and all the experiments are scripted in the MATLAB software package (2016b, MathWorks, USA).

### Behavior Training

All animals were trained in a color discrimination experiment based on a Go/NoGo paradigm. To motivate task acquisition, we used water as a reinforcement. Pigeons were water-restricted for the whole night before the experiment. All pigeons learned to mandibulate (moving the lower jaw) during the reward-associated color presentation (Go stimulus) and to inhibit their response to a non-rewarded color (NoGo stimulus). Pigeons were randomly assigned into two groups to counterbalance the associated color with reward. For one group, the green color was associated with reward and served as a Go stimulus and the green color as a NoGo stimulus (S-, no response), while for the other group the colors were reversed. In addition, to avoid animals’ attention shifting to the light intensities instead of the light colors, we used two different light intensities for both colors (20% or 100% of the maximum light intensity 28.9 cd/m2, respectively) during the experiment. The results indicated no behavioral difference in response to different intensities (paired-sample *t*-test, *p*= 0.69 (Go trials), *p*= 0.62 (for NoGo trials)) during the conditioning phase.

The behavioral training steps have already been explained in the study by Behroozi et al.^29^ and is based in its operant components on a method introduced by Mallin and Delius^101^. In summary, after habituating animals to the head restrainer, they were trained for an operant conditioning experiment based on a Go/NoGo paradigm. The training steps consisted of autoshaping and instrumental conditioning phases. In the first phase of the autoshaping, the S_+_ (red/green light as a conditioned stimulus for the corresponding group) was presented for 2 s which was followed by water access irrespective of the animals’ response (classical conditioning). The water reward was signaled using two white lights (reward light) and followed by a randomly jittered 12.2-20.2 s ITI. After reaching the behavioral performance of 80%, the experimental phase was shifted to the instrumental conditioning procedure. In this phase, pigeon’s mandibulations during the S+ have only resulted in immediate water access. When pigeons’ behavioral performance reached 70% correct responses, the third phase of autoshaping was started. In the third phase, the pigeons’ correct responses did not result in an immediate reward. The reward was delivered 800ms after the CS+ offset. The main experiment started when the behavioral performance reached 80%. The training was followed by adding a NoGo stimulus (non-conditioned stimuli; S_-_) to the experimental procedure. Pigeons underwent a Go/NoGo task training and learned to operantly mandibulate upon seeing S+ and to inhibit their response upon seeing an S-. Mandibulation during S-presentation did not result in any reward or punishment. Main fMRI data for the conditioning phase was collected when animals mandibulated correctly in at least 80 % of the trials on 2 consecutive training days. Each fMRI run consisted of two blocks, each containing 72 trials (36 Go and 36 NoGo). The experiment included three resting-state scans: an initial resting-state scan before block 1, an intermediate resting-state scan between the two blocks, and a final resting-state scan after block 2. The initial and final resting-state scans lasted 10 minutes each, while the intermediate resting-state scan lasted 20 minutes. Twenty-four hours after the conditioning, pigeons entered the extinction phase. The task procedure for the extinction phase was the same as the conditioning phase, except that mandibulation during S+ presentation was not rewarded, and water was not delivered at any time.

### fMRI acquisition

Neuroimaging data was acquired using a Bruker BioSpec small animal MRI scanner system (7T horizontal bore, 70/30 USR, Germany) equipped with an 80 mm quadrature birdcage resonator for radiofrequency transmission, a single-loop 20-mm surface coil for signal detection, and ParaVision 5.1 software interface.

The imaging sequences began with the acquisition of three runs (horizontal, coronal, and sagittal) of the scout-set, using Rapid Acquisition with Relaxation Enhancement (RARE) sequence. Scan parameters were as follows: TR = 4s, TE_eff_ = 40.37 ms, RARE factor = 8, no average, acquisition matrix = 256 × 128, FOV = 32 × 32 mm, spatial resolution = 0.125 × 0.25 mm2, slice thickness = 1 mm, number of slices = 20 horizontal, 17 sagittal, and 15 coronals. These series of scout images were used to localize the position of the animal’s brain to place 11 coronal slices to record functional images that cover the entire telencephalon.

Functional images were acquired using a single-shot RARE sequence^29^. Scan parameters were as follows: TR = 4000 ms, TE_eff_ = 41.58 ms, partial Fourier transform accelerator = 1.53, encoding matrix = 64 × 42, acquisition matrix = 64 × 64, FOV = 30 × 30 mm^2^, in-plane spatial resolution = 0.47 × 0.47 mm^2^, radio-frequency pulse flip angles for excitation and refocusing = 90°/180°, slice thickness = 1 mm, no slice distance, slice order = interleaved, excitation and refocusing pulse form = scanner vendor gauss512, receiver bandwidth = 50,000 Hz. Subsequently, two saturation slices were used to avoid artifacts caused by eye movements by suppressing signals from the eyes.

In-plane anatomical images with the same orientation and position as the functional slices were acquired. Scan parameters were as follows: TR = 4000 ms, TE = 36.52 ms, RARE factor = 8, FOV= 30 × 30 mm^2^, acquisition matrix = 128 × 128, in-plane spatial resolution = 0.23 × 0.23 mm^2^, number of slices = 34, slice thickness = 0.25 mm. These structural scans were used for enhancing the registration of lower-resolution functional images to the higher resolution anatomical images.

High-resolution T2-weighted anatomical images were acquired using a RARE sequence. Scan parameters were as follows: TR = 2000 ms, effective TE = 50.72 ms, RARE factor = 16, number of averages = 1, FOV = 25 × 25 × 15 mm^3^, Matrix size = 128 × 128 × 64, spatial resolution= 0.2 × 0.2 × 0.23 mm^3^.

## Data Analysis

### Behavioral Data Analysis

Behavioral analyses were performed using MATLAB (The Mathworks, Natick, MA) and SPSS 21 (IBM Corporation, Armonk, NY, USA). The timestamped occurrence of mandibulations registered by the piezo was used as a behavioral readout. The main dependent variable was the fractional mandibulation rate, defined as the percentage of trials in which an operant response occurred. For each trial, if at least one response detected during the stimulus-on period, the fractional response for that trial was recorded as 1; otherwise, it was recorded as 0. Mean responses for S+ and S+ were calculated for the conditioning and extinction phases as well as for the early and late phases of the extinction phase with the early and late phases corresponding to the first and second blocks, respectively.

Behavioral data were then analyzed for the conditioning and extinction phases separately. For the conditioning phase, the data were analyzed using a paired t-test. For the extinction phase, the data were analyzed using a paired t-test for the entire extinction phase (block 1 + block 2) and a two-way ANOVA was used to determine the effects of stimulus type (S+ vs. S+ ; first independent variable) and extinction phase (early vs. late; second independent variable) on fractional responses. Appropriate post hoc t-tests were conducted to analyze significant effects further. Additionally, to illustrate the gradual decline in responding during extinction in greater detail, fractional responses were grouped into bins of six consecutive trials for both S+ and S- and plotted.

### fMRI data analysis

Preprocessing and statistical analyses were carried out using FSL version 6.0.6.5 (www.fmrib.ox.ac.uk/fsl)^102^ and custom-made scripts in MATLAB (Mathworks, Natick, MA, USA).

The following preprocessing steps have been applied to minimize noise fluctuations and improve the detection of activation: upscaling the voxel size by a factor of 10, discarding the first 10 volumes to ensure longitudinal magnetization reached steady-state, motion correction, slice time correction, non-brain matter removal from functional data using a brain mask, spatial smoothing (8 mm FWHM Gaussian kernel) to increase the signal-to-noise ratio, global intensity normalization by a single multiplicative factor, and high-pass temporal filtering (cut-off period 90 s).

To enhance registration, functional images were co-registered to in-plane anatomical images using affine linear registration (six degrees of freedom), then to the high-resolution T2-weighted RARE anatomical images using affine linear registration (12 degrees of freedom), and finally to a population-based template generated based on the mean 3D high-resolution anatomical images of all subjects using linear registration with FSL FLIRT.

FSL FEAT package was used for statistical analysis. A general linear model (GLM) was created for the Go/NoGo task with each subject’s GLM included 10 regressors. The first four regressors were modeled separately for each block: Go trial with mandibulation, Go trial with no mandibulation, NoGo trial with mandibulation, and NoGo trial with no mandibulation. Additionally, two separate regressors were included for mandibulation: one for mandibulation occurring within 5 seconds after stimulus offset and one for mandibulation during rest periods and inter-trial intervals (ITIs), excluding the post-stimulus period. These two regressors were modeled across the entire experiment rather than per block. All Regressors were modeled through convolution with the estimated HRF in the pigeon brain^29^.

### Contrasts for first-level analysis

Go trials without mandibulation during the stimulus-on period were used to characterize the successful extinction learning effect (S+_ext_), while NoGo trials without mandibulation during the stimulus-on period served as a control (S-_ext_). For each block, S+_ext_ > S-_ext_ contrasts were created separately to differentiate between early and late effects^103,104^. Temporal derivatives were included in the model to account for variability in the timing of the HRF across brain regions and subjects. Besides, motion outliers assessed with the FSL build-in function (fsl_motion_outlier) were added as confounds to account for any residual effects due to movements.

### Contrasts for higher-level analysis

The two contrasts of interest for the early and late phases, S+_extE_ > S-_extE_ and S+_extL_ > S-_extL_, were carried forward to the higher-level analysis using the mixed-effect model (FLAME1 + 2). All fMRI data shown were cluster-corrected for multiple comparisons at z = 3.1, p < 0.05.

Group results were co-registered linearly to the pigeon MRI atlas and 3D MRI images and were visualized using the Mango software (http://ric.uthscsa.edu/mango/, version 4.1). Anatomical localization within each cluster was obtained by searching anatomical borders based on the pigeon MRI atlas Güntürkün et al.^52^.

## Author contributions

M.B. and O.G. designed the study. M.B., M.G. and X.H. conducted the fMRI experiment. A.S., M.G., E.G. and M.B participated in data processing and analysis. M.B. and O.G. supervised the work. A.S., M.B., and O.G. wrote the original draft. O.G. contributed to funding acquisition and resources. All authors contributed to the manuscript review and editing process.

## Data and code availability statement

Related data processing codes are available at https://github.com/mehdibehroozi/pigeon_fMRI.

## Acknowledgments

This work was supported by the Deutsche Forschungsgemeinschaft (DFG, German Research Foundation) through grant SFB 1280 (A01, A03, F01, F02) project number 316803389 and AVIAN MIND, ERC-2020-ADG, LS5, GA No. 101021354.

